# Universality of phenotypic distributions in bacteria

**DOI:** 10.1101/2022.08.21.504683

**Authors:** Kuheli Biswas, Naama Brenner

## Abstract

Some phenotypic properties in bacteria exhibit universal statistics, with distributions collapsing under scaling. The extent and origins of such universality are not well understood. Using phenomenological modeling of growth and division, we identify compound “shape-factors” that describe the distributions throughout a large set of single-cell data. We find that the emergence of universal distributions is associated with the robustness of shape-factors across conditions, explaining the universality of cell size and highly expressed protein content and the non-universality of times between consecutive divisions. A wide range of experimental data sets support our theory quantitatively.

---

Bacterial growth and division have fascinated scientists and have been studied for decades. Mathematical models to account for the variability in cell size and division times across time and among individuals were developed early on [1–4], but the comparison to data at the time was limited. More recent single-cell imaging and microfluidics technology yields high-quality data with large statistical power [5–12], enabling to test quantitative models of growth and division and the relation between temporal dynamics and statistical properties.

Previous works have reported that the distributions of cell size and highly expressed proteins are non-Gaussian skewed, well approximated by a log-normal form [7, 13– 16]. These distributions have a universal shape, as manifested by their collapse under scaling [17, 18]. In physics, such distribution collapse is associated with the insensitivity of macroscopic properties to underlying microscopic ones [19]. In biological systems, analogous collapse appears in different contexts, but its origin and extent are still not well understood [20**?**, 21]

The distribution of times between divisions has also been extensively studied, but here the emerging picture is less conclusive. Some experiments report a Gaussian-like distribution of division times [8, 9, 18, 22] while others find more skewed distributions [1, 2, 12, 23–25]. Some of these report a universal shape for division times [26, 27]. Modeling work assuming a small-noise approximation has concluded that times between divisions are distributed approximately Gaussian [15]. In contrast, “sloppy size control” models [28] as well as other modeling approaches [29–31] find skewed distributions. Thus, a unifying framework that can explain the observed diversity and apparent disagreement in division time distribution shapes is lacking.

Here, we compute the division time distribution using *first passage time* techniques in a phenomenological coarse-grained stochastic model of growth and division. We find that the ratio between two stochastic parameters - variability in exponential growth rate and in threshold - defines a dimensionless “shape-factor” that governs the distribution shape beyond shift and scale transformations. Analyzing a large set of single-cell data from microfluidic experiments, we find that division time dis-tributions span a range of diverse behaviors, from almost Gaussian to very skewed. Our theoretical predictions, based on direct estimation of dynamic parameters, agree with the data throughout this range of behaviors. Combining these observations with known properties of cell size and highly expressed protein content, we address the question of universality. By observing the spread of data points in the landscape of shape-factors, we find that for cell size and protein, the data adhere near a manifold of fixed value, while for division times they scatter through multiple contour lines. This finding sheds light on the universality of size and protein distributions, and the non-universality (or shape diversity) of division time distributions. It highlights the emergence of compound parameter combinations, whose sensitivity is key to understanding the system’s behavior.

## Model Definition

High-resolution single-cell measurements have revealed that several bacterial species, including *E. coli*. grow smoothly and exponentially across each cycle [8, 24, 25]; division occurs abruptly - consistent with the idea of triggering by some threshold event (Fig. 1 a). The growth dynamics during the *n*^*th*^ cycle can thus be approximated as

**FIG. 1.**
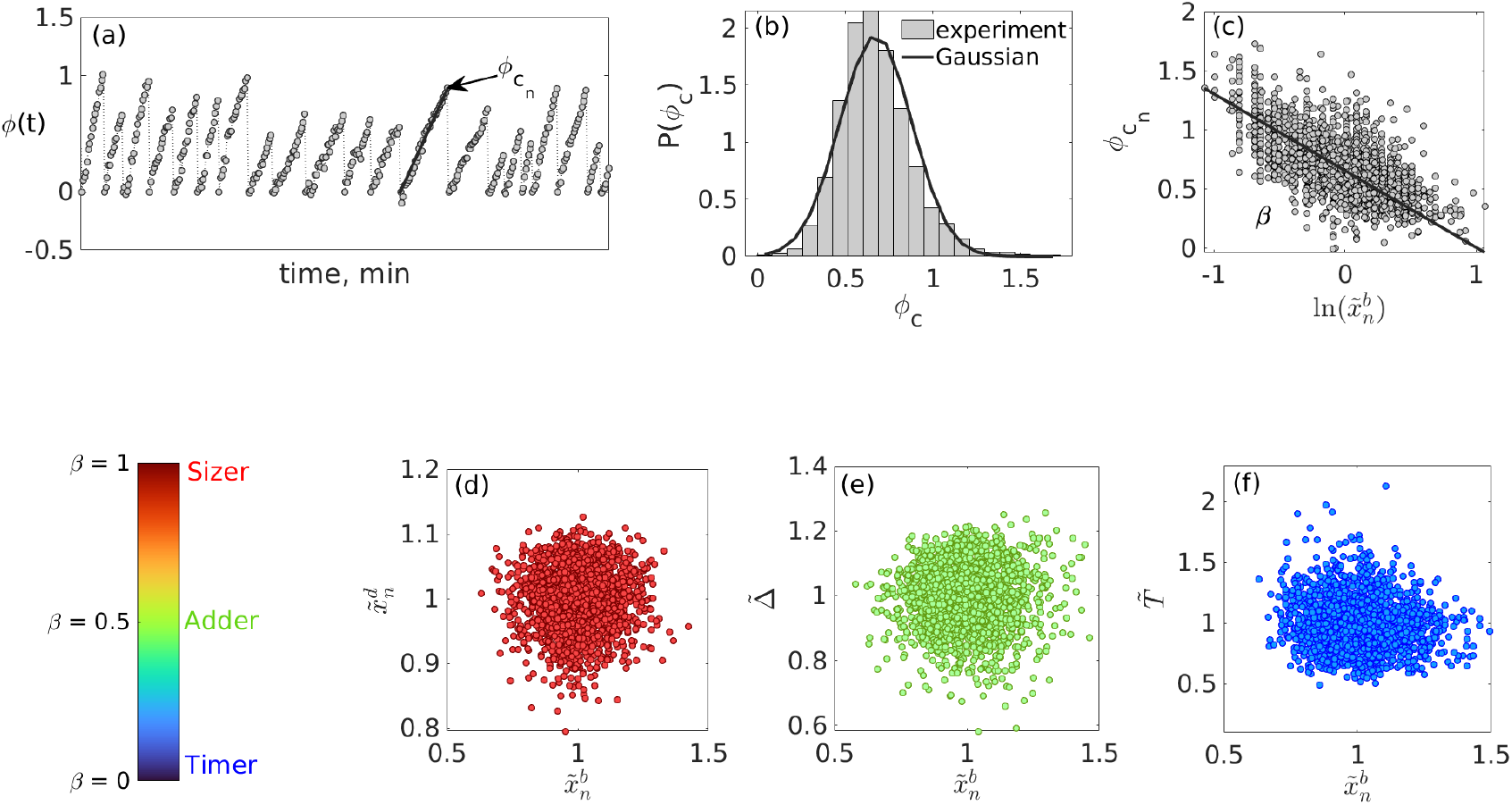
Phenomenological model unifying different division control mechanism. **(a-d)**: Data from [25], where *E. coli* were grown in a microfluidic device. **(a)** In exponential growth, Logarithmic Fold Change (LFC) in cell size, *φ*(*t*) = *αt*, is linear across cycles. **(b)** At division, LFC reaches a stochastic threshold *φ*_*c*_, with distribution *P* (*φ*_*c*_) approximately Gaussian (*µ*_*φ*_ ≈ ln 2, *σ*_*φ*_ = 0.17). **(c)** Correlations in *φ*_*c*_ across consecutive cycles are negligible with correlation coefficient ≈ -0.02. **(d)** The threshold *φ*_*c*_ is negatively correlated with normalized birth size 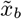 with best-fit slope *β* = 0.65. Experimental details are provided in [32]. **(e-g)** Scatterplots from simulations of the phenomenological model, Eqs. 1, 2: division size (d), added size (e) and cycle time (f) vs. normalized initial size. The independence of these variables on initial condition is often interpreted as indicating a control mechanism where they are kept constant. By tuning the homeostasis parameter *β*, the model reduces to previously studied modes of division control: **(d)** sizer (*β* = 0.9), **(e)** adder (*β* = 0.5), and **(f)** timer (*β* = 0.2) as special cases. 10^4^ generations were simulated with *α* = 0.023 min^−1^ and *ξ* = *𝒩* (0, 0.15). See [32] for more details.

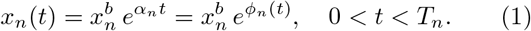

Here *x*_*n*_ is the cell size or highly expressed protein content, with 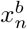 the value at birth. *α*_*n*_ is the effective growth rate, and *φ*_*n*_(*t*) the Logarithmic Fold Change (LFC) in cell size. The cycle ends at time *t* = *T*_*n*_ with symmetric division. We formulate a generalized stochastic threshold-crossing model of division that unifies different previously studied division mechanisms as special cases.

The LFC along the *n*^*th*^ cycle is by definition zero at birth *φ*_*n*_(0) = 0, and at division reaches a fluctuating values 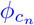 (Fig. 1a). These fluctuations are well described by a Gaussian distribution around ln 2 (Fig. 1b), with negligible correlations across cycles (Fig. 1c) and a significant negative correlation with cell size at birth (Fig. 1d). These statistical properties are all captured by the following phenomenological expression [24]

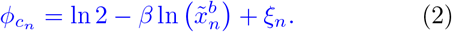

Here *ξ*_*n*_ is a Gaussian white noise with zero mean, 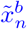 is the birth size normalized by its mean, and 0 *< β* ≤ 1 is the slope of Fig. 1d. Due to the lognormal distribution of size, the correlation term does not modify the Gaussian nature of *φ*_*c*_; at the same time, such a negative correlation induces a stable size distribution over multiple generations, interpolating between previously described division models [15, 16, 24, 33, 34]. To demonstrate this, we simulate the model Eqs. (1, 2) and present the resulting scatterplots of different variables vs. initial cell size in Fig. 1(e-g). Such plots have been extensively used to identify the mode of division control. For example in Fig. 1f, *β* = 0.5 and added size is independent of the birth size, a correlation known as the “adder” mode of division control. Sizer and timer-like correlations are found for other values of *β* in Fig. 1(e,g). The mathematical mapping between the current and previously formulated models is elaborated in [32].

## Model Solution

Eqs. (1, 2) together constitute a phenomenological model that was successfully solved for cell size and protein distributions [15, 16], and will be analyzed below in terms of division time statistics. We assume that cell division occurs when the LFC crosses a fluctuating process; this does not necessarily imply a mechanistic interpretation, but rather a tool for calculation. Using the first passage time (FPT) technique [35, 36]. Cell division occurs when the Gaussian process *P* (*φ*_*c*_) = 𝒩 (ln 2, *σ*_*φ*_) crosses the absorbing boundary *φ*(*t*) = *αt* [37]. The survival probability - the probability that the cell has not divided till time *t* - can then be expressed as

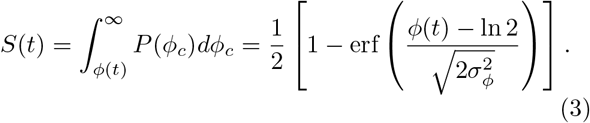

Fo r a fixed growth rate *α*, this is related to the probability density of the FPT by

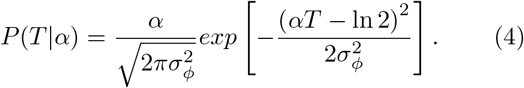

However, the growth rate estimated from the slope of *φ*(*t*) varies across cycles and follows Gaussian distribution, *P* (*α*) = 𝒩 (*µ*_*α*_, *σ*_*α*_) (Fig. 2a). To account for variable growth rates, we integrate Eq. (4) with *P* (*α*), which leads to the following division time distribution [32]

**FIG. 2.**
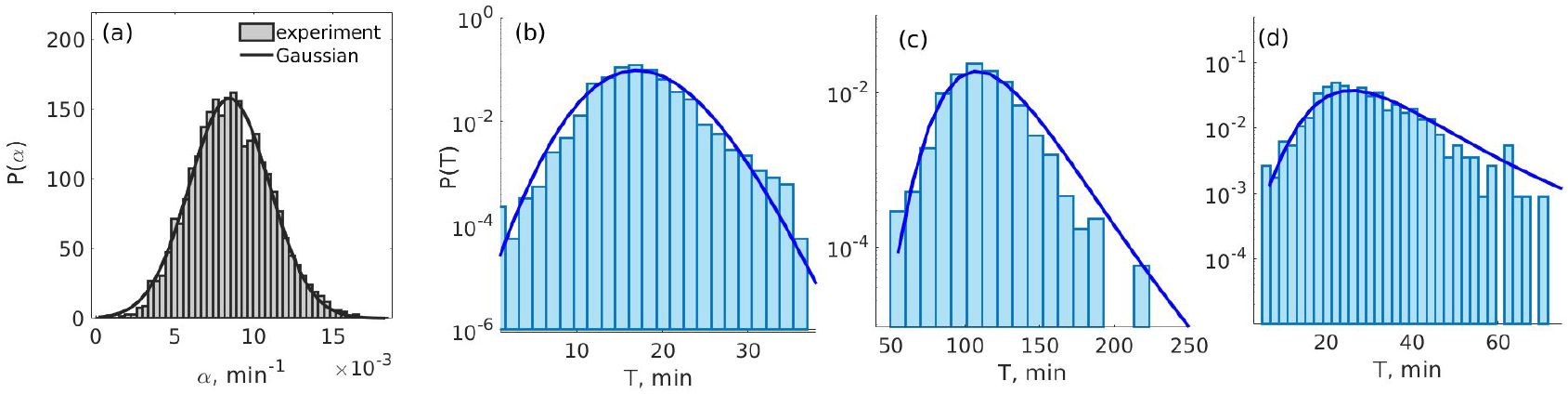
Division time distributions are predicted across a range of shape-factors. **(a)** Growth rates *α*_*n*_, obtained from the slope of *φ*_*n*_(*t*) (Fig. 1a) follow a Gaussian distribution, *P* (*α*) = 𝒩 (*µ*_*α*_, *σ*_*α*_). These data from [25] show *χ*_*α*_ = *σ*_*α*_*/µ*_*α*_ = 0.27. **(b)-(d)** Division time distributions from data (bars) corresponding to different Σ_*T*_ are well predicted by Eq. (5) (solid lines). The shape factors are (b) Σ_*T*_ = 0.64 [8], glucose medium, (c) Σ_*T*_ = 1.3 [11] arginine medium, (d) Σ_*T*_ = 1.8 [24], LAC-M9 medium. In all experiments, *E. coli* bacteria were grown in mother machines; see details in [32].

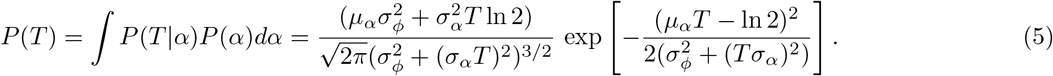

Transforming this expression to dimensionless time [32], we find that two dimensionless parameters appear in the formula for division times: *σ*_*φ*_, a measure of threshold variance, and *χ*_*α*_ = *σ*_*α*_/*µ*_*α*_, a measure of growth-rate variability. The shape of the distribution is strongly influenced by the ratio of these two parameters, which reflects the relative strength of the two noise sources in the model: Σ_*T*_ = *χ*_*α*_/*σ*_*φ*_. We term this ratio the shape-factor; it defines regions where the distribution resembles a Gaussian shape when Σ_*T*_ *<* 1, and a skewed heavy-tailed shape when Σ_*T*_ *>* 1.

Fig. 2(b-d) shows a few examples taken from different experiments, illustrating a range of shape factors from Σ_*T*_ = 0.64, an almost Gaussian shape, to a larger value of Σ_*T*_ = 1.8, where the shape is highly skewed. The experimental data (bars) are well described by Eq. 5 (lines) throughout this range of behaviors. A larger collection of division time distributions can be seen in [32], all in good agreement with the theory. Note that there are no fitting parameters in this theory. To obtain the theoretical lines, we estimate stochastic variables *µ*_*α*_, *σ*_*α*_ and *σ*_*φ*_ directly from dynamic trajectories such as those in Fig. 1(a). These are then inserted into Eq. (5).

## Universal and non-universal distributions

The stochastic LFC threshold model was used in previous work to predict a log-normal distribution of cell size and protein content; its shape depends on the compound shape-factor 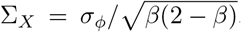, which in turn depends on the threshold standard deviation *σ*_*φ*_ and the homeostasis parameter *β* [15, 16]. Here we identified the compound parameter Σ_*T*_ = *σ*_*φ*_/*χ*_*α*_ as determining the shape of the division time distribution. In contrast to the broad range of distribution shapes found for division time, cell size and highly expressed protein content have been consistently reported to exhibit a universal distribution shape[14, 15, 17, 18, 27]. We next investigate properties of the shape-factors to shed light on the question of universal distribution collapse.

In Fig. 3(a,b) we assemble data from a large set of single-cell experiments in the planes of stochastic parameters (*χ*_*α*_, *σ*_*φ*_) and (*β, σ*_*φ*_) respectively. Each point represents one experiment, where single-cell traces were pooled for estimation. In all cases, bacteria were grown in mother machines, but relevant conditions such as medium, temperature, or bacterial strain vary significantly among experiments (see Table 1 in [32]). In addition, in these parameter planes, the shape-factor for division time (a) and cell size (b) are depicted as grey contour lines of constant values. Examining the embedding of the data across the shape landscape reveals an important difference between the cases.

**FIG. 3.**
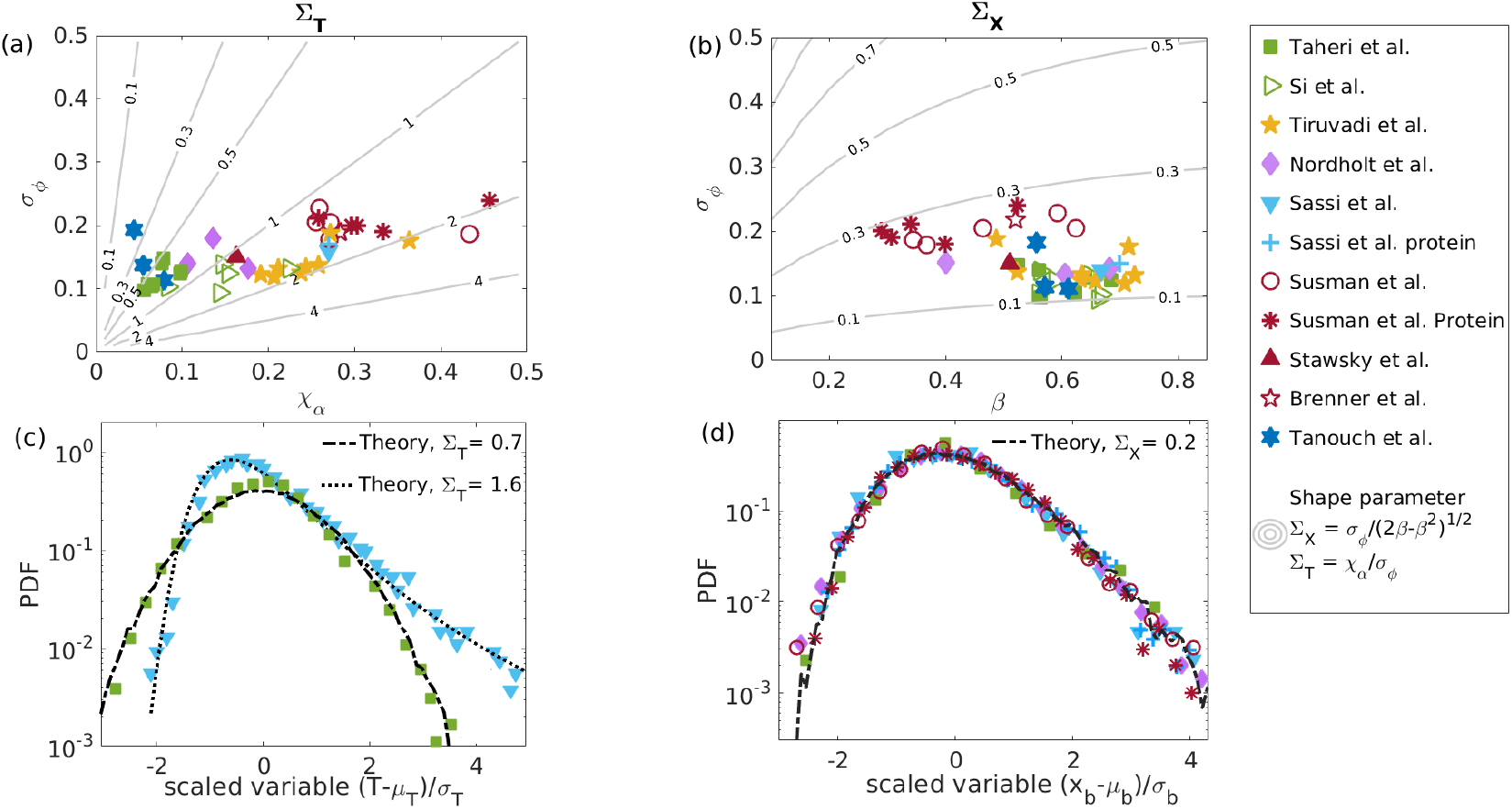
Non-universal division time distribution and universal cell size distribution. **(a)** Plot of *χ*_*α*_ vs *σ*_*φ*_, estimated from experimental data, span a broad range of shape factors Σ_*T*_ (gray contours). **(b)** Plot of *σ*_*φ*_ vs. *β*, estimated for the same data, cluster around a small range of shape factor for cell size or protein Σ_*X*_ (gray contours). **(c)** The scaled distribution of cell division times is non-universal, showing a Gaussian-like distribution for Σ_*T*_ *<* 1 (green squares, [8]) and a skewed shape for Σ_*T*_ *>* 1 (blue triangles, [25]). The black curves are obtained from our theory Eq. (5). **(d)** The scaled distribution of cell size and protein number collapse under two-parameter scaling ([8, 22, 24, 25]), and agree well with log-normal distribution [16] (black curve). Data are taken from [7–12, 22, 24, 25, 38] and experimental details are provided in Table 1 of [32].

In (a), the data scatter across many contour lines, spanning a broad range of shape-factor values (Σ_*T*_ ∼ 0.1 to 3). Contributing to this variability is the broad range of growth rate noise *χ*_*α*_ (x-axis), approximately ten-fold 0.05 to 0.5, to which the shape-factor is inversely proportional. This is consistent with the broad range of behaviors we observed for division time distributions across data sets. Two examples of scaled distributions are shown in panel (c). In (b), in marked contrast, the data are concentrated in a small region around a single contour line, maintaining an almost constant shape-factor for cell size (Σ_*X*_ ≈ 0.2). It has previously been demonstrated that the shape of the lognormal distribution is very weakly sensitive to the shape-parameter value in this regime [16]; together these observations are consistent with the collapse of these distributions upon scaling (panel (d)).

## Discussion

We examined single-cell statistical properties using a phenomenological growth and division model. This model relies on several key assumptions: exponential growth during the cell cycle, variable exponential rates in successive cycles, and symmetric division triggered by a noisy threshold in Log Fold Change of cell size - negatively correlated with birth size. This approach has been validated in previous studies, and encompasses various homeostasis modes, including adder, sizer, and timer scenarios, as specific instances. [39]

By employing a first passage time framework, we computed the distribution of division times. The approximations made in obtaining this result do not hinge on a small-noise approximation in growth rate across cell cycles (see [32]). In the data we analyzed, encompassing a wide spectrum of experiments in microfluidic traps, the small-noise limit only applies to a minority of experiments. The range of behaviors - from very narrow to very broad growth rate distribution - served as a central motivation for extending the phenomenological theory in our present study.

The resulting distribution has strictly no finite moments, but a limited range of data allows for a good comparison with an appropriate cutoff. Notably, this result accurately predicts the relationships between data moments and model parameters, as detailed in [32]. The long tails of the distribution result from the Gaussian assumption for growth rates. Alternative assumptions, such as a Gamma distribution, yield empirically similar distributions [32], but do not provide an analytic expression that allows for identifying parameter dependencies and a clearly understandable shape-factor.

The shape-factor Σ_*T*_ governs the distribution shape beyond shift and scale, eliminated by two-parameter scaling. An analogous factor had previously been found for cell size, but its empirical values and significance for distribution scaling were not studied. By comparing the two shape-factors we were able to illuminate the sources of distribution universality, as stemming from the robustness of these parameter combinations as experimental conditions are varied. This explanation remains phenomenological and does not clarify the source of difference in robustness. In particular, the reasons behind different levels of variability in single-cell growth rates among experimental conditions remain to be understood. It can be speculated that the sensitivity of growth rates to their environment and their strong correlation with division times over long timescales [38] make them effective control variables that require flexibility for compensation. On the other hand, cell size does not partake in long-term homeostatic correlations; stable growth can occur at various cell sizes, rendering it an “irrelevant variable” in the sense of statistical mechanics with a universal distribution. These hypotheses warrant further investigation.

The description of cell size and highly expressed proteins has converged to a similar modeling framework. Both quantities adhere to the fundamental model assumptions outlined above and exhibit strong cycle-by-cycle correlations between their stochastic variables [7, 24, 25]. Consequently, in Fig. 3 we found that they share the same universal distribution shape. Intriguingly, the statistics of division time can be predicted based on the dynamics of either cell size or any highly expressed protein, with similar predictive success [32]. Thus, while cell division is likely triggered by multiple events, their tight interconnection reduces the complexity of the problem, allowing several options to predict statistical properties effectively. This aligns with previous observations of balanced exponential growth as a dynamic attractor characterized by robust relationships among constituents [31]. Understanding such reduction in effective dimensionality in high-dimensional yet highly correlated systems like the biological cell remains an important open question.

## Acknowledgements

K.B. is supported by a fellowship from the Lady Davis Foundation at the Technion. The research is supported in part by the Israeli Science Foundation (grant number 155/18), and the Binational Science Foundation (grant number 2016376). We are grateful to Erez Braun and Ron Teichner for comments on the manuscript.

## AI declaration

While preparing this work, the authors used ChatGPT to improve the readability and language of the work. After using this tool, we reviewed and edited the content as needed and take full responsibility for the paper’s content.

